# Lysosomal positioning regulates Rab10 phosphorylation at LRRK2-positive lysosomes

**DOI:** 10.1101/2020.12.01.406223

**Authors:** Jillian H. Kluss, Alexandra Beilina, Chad D. Williamson, Patrick A. Lewis, Mark R. Cookson, Luis Bonet-Ponce

**Author notes:** Correspondence: Luis Bonet-Ponce, Ph.D., Cell Biology and Gene Expression Section, Laboratory of Neurogenetics, National Institute on Aging, NIH, 35 Convent Drive, Room 1A–108, Bethesda, MD, 20892–3707, USA. Correspondence: Mark R. Cookson, Ph.D., Cell Biology and Gene Expression Section, Laboratory of Neurogenetics, National Institute on Aging, NIH, 35 Convent Drive, Room 1A–116, Bethesda, MD, 20892–3707, USA.

## Abstract

Genetic variation at the *Leucine-rich repeat kinase 2 (LRRK2)* locus contributes to enhanced lifetime risk of familial and sporadic Parkinson’s disease. Previous data have demonstrated that recruitment to various membranes of the endolysosomal system results in LRRK2 activation. However, the mechanism(s) underlying LRRK2 activation at endolysosomal membranes and the cellular consequences of these events are still poorly understood. Here, we directed LRRK2 to lysosomes and early endosomes, triggering both LRRK2 autophosphorylation and phosphorylation of the direct LRRK2 substrates Rab10 and Rab12. However, when directed to the lysosomal membrane, pRab10 was restricted to perinuclear lysosomes, whereas pRab12 was visualized on both peripheral and perinuclear LRRK2-positive lysosomes, suggesting that lysosomal positioning provides additional regulation of LRRK2-dependent Rab phosphorylation. Anterograde transport of lysosomes to the cell periphery by increasing expression of ARL8B and SKIP or by knockdown of the motor adaptor protein JIP4 blocked recruitment and phosphorylation of Rab10 by LRRK2. Conversely, overexpression of the Rab7 effector protein RILP resulted in lysosomal clustering within the perinuclear area and increased LRRK2-dependent Rab10 recruitment and phosphorylation. The regulation of Rab10 phosphorylation in the perinuclear area depends on counteracting phosphatases, as knockdown of phosphatase PPM1H significantly increased pRab10 signal and lysosomal tubulation in the perinuclear region. Our novel findings suggest LRRK2 can be activated at multiple cellular membranes including lysosomes, and that lysosomal positioning further provides regulation of some Rab substrates likely via differential phosphatase activity in nearby cellular compartments.

## INTRODUCTION

Coding mutations in the *LRRK2* gene can cause familial Parkinson’s Disease (PD) (Paisán-Ruíz et al. 2004; Zimprich et al. 2004), and non-coding variants in the promoter of the same gene act as risk factors for sporadic PD (Nalls et al. 2019). *LRRK2* encodes Leucine-rich repeat kinase 2 (LRRK2), a large protein consisting of dual enzymatic domains flanked by protein-protein interaction scaffold domains (Roosen and Cookson 2016). Known LRRK2 pathogenic mutations are located in the ROC/GTPase, COR and kinase domains, and produce a toxic hyperactive protein (Greggio et al. 2006; West et al. 2005), implying that kinase inhibitors reducing LRRK2 activity may be useful to treat PD. Therefore, defining the mechanism(s) that regulate LRRK2 activity is of critical importance for understanding both disease risk and potential therapeutics targeting LRRK2.

LRRK2 phosphorylates itself at Serine 1292 (Sheng et al. 2012) as well as Serine/Threonine residues in a conserved region of the switch II domain of a subset of Rab GTPases, leading to differential interactions with effector proteins and thus linking LRRK2 to vesicle mediated transport (Steger et al. 2016; Bonet-Ponce and Cookson 2019). Likely due to the phosphorylation of Rab GTPases, LRRK2 has been nominated to play important roles at endomembranes including early endosomes, recycling endosomes, late endosomes, and lysosomes (Roosen and Cookson 2016; Hur et al. 2019; Yun et al. 2015; Bonet-Ponce and Cookson 2021). We and others have previously shown that damage to membranes within the endolysosomal system results in LRRK2 activation as measured by Rab phosphorylation (Bonet-Ponce et al. 2020; Herbst et al. 2020; Eguchi et al. 2018). LRRK2-dependent Rab phosphorylation at lysosomes results in recruitment of JNK interacting protein 4 (JIP4), a motor adaptor protein, and formation of tubules from the lysosomal surface (Bonet-Ponce et al. 2020). LRRK2 may also play an important role in regulation of lysosomes *in vivo,* as lysosomal proteins are dysregulated in LRRK2 knockout mice (Pellegrini et al. 2018) or after chronic LRRK2 kinase inhibition (Kluss et al. 2021). Intriguingly, the same treatments result in variable effects on Rab phosphorylation with Rab12 being strongly inhibited compared to Rab10 and Rab29 *in vivo* (Kluss et al. 2021).

Although these prior data imply that LRRK2 is part of the cellular response to membrane damage, the mechanism(s) by which LRRK2 becomes activated at membranes are not fully understood. A prior study using a mitochondrially-tagged Rab29 protein suggested that membrane identity is unimportant in LRRK2 activation, as directing Rab29 to mitochondria is sufficient to activate LRRK2 and results in Rab10 phosphorylation (Gomez et al. 2019). However, whether similar events occur at lysosomes (or other compartments), which have additional regulatory signals including positioning (Ballabio and Bonifacino 2020), is currently not defined.

Here, we designed two independent methods to deliver LRRK2 to early endosomes and lysosomes in order to investigate potential membrane-dependent patterns of activation of LRRK2 and its direct substrates Rab10 and Rab12 at sites T73 and S106, respectively. We found that LRRK2 becomes kinase active at either membrane, as indicated by autophosphorylation at site S1292 and was able to recruit and phosphorylate both Rab10 and Rab12. However, further analysis showed that only a subset of perinuclear LRRK2-positive lysosomes could recruit pRab10, whereas pRab12 was shown at almost all LRRK2-positive lysosomes throughout the cell. To address whether spatial location is important for the recruitment of pRab10 to lysosomes, we directed lysosomes to the cell periphery via ARL8B (ADP Ribosylation Factor Like GTPase 8B) and SKIP (SifA and kinesin-interacting protein) overexpression or JIP4 siRNA knockdown. Additionally, we directed lysosomes to the juxtanuclear area via LLOMe (L-leucyl-L-leucine methyl ester) treatment or RILP (Rab7A-interacting lysosomal protein) overexpression. We found that peripherally targeted LRRK2-positive lysosomes were unable to recruit pRab10, and that clustering lysosomes into the perinuclear area significantly increased pRab10 signal and colocalization. Interestingly, pRab12 was not sensitive to lysosomal position, suggesting that additional signaling events may control the phosphorylation of specific Rabs on subsets of lysosomes. To address a possible mechanism for this observation, we knocked down PPM1H (Protein Phosphatase, Mg2+/Mn2+ Dependent 1H), a phosphatase known to dephosphorylate T73 Rab10 that is reported to be associated with the Golgi apparatus and thus modulates LRRK2 signaling (Berndsen et al. 2019). PPM1H knockdown resulted in significant increase in pRab10 presence as well as frequency of pRab10/JIP4-positive lysosomal tubules. These novel findings suggest LRRK2-dependent recruitment and subsequent phosphorylation of Rab10 at lysosomes is in part affected by lysosome position, and that PPM1H plays a role as a limiting factor in pRab10-dependent tubulation at perinuclear lysosomes. This mechanism may be of particular importance when elucidating the LRRK2-specific mechanisms of PD pathogenesis.

## RESULTS

### LRRK2 is activated at early endosomal or lysosomal membranes using two orthogonal methods of relocalization

LRRK2 has been reported to play a role in various compartments of the endolysosomal system (Bonet-Ponce and Cookson 2021). To study the activation of LRRK2 in this context, we designed two orthogonal methods to direct LRRK2 to lysosomal or early endosomal membranes. We chose these two distinct membranes as they are spatially and functionally distinct portions of the endolysosomal pathway. First, we took advantage of the FKBP-FRB system using rapamycin-binding domains from FKBP12 and mTOR, respectively. In the presence of rapamycin, these domains can form a heterodimer, and rapidly and irreversibly direct a target protein to a target membrane (Robinson, Sahlender, and Foster 2010). We therefore tagged LRRK2 with the FKBP sequence (3xFLAG-FKBP-LRRK2), and fused the FRB sequence to the lysosomal and early endosomal membrane markers, LAMP1 and Rab5 respectively. Secondly, we cloned two new chimera-LRRK2 constructs by tagging the N-terminus of LRRK2 with transmembrane domains specific to lysosomes and early endosomes from LAMTOR1 and HRS (FYVE finger protein localized to early endosomes), respectively (Raiborg et al. 2001; Nada et al. 2009).

We first confirmed the efficiency of translocation of LRRK2 to the lysosomal membrane. Co-transfection of the FKBP-LRRK2 and LAMP1-FRB-CFP constructs into HEK293FT cells resulted in strong colocalization of LRRK2 to the LAMP1 construct in the presence of rapamycin (Fig 1A). Similarly, the LAMTOR1(1-39aa)-LRRK2 chimera (referred to as LYSO-LRRK2) colocalized with endogenous LAMP1, LAMTOR4, and CTSD (Fig 1A and Supplementary Fig 1A-C). Having established correct localization of our constructs, we next evaluated whether the direction of LRRK2 to lysosomes was sufficient to activate the kinase activity of LRRK2. Cells transfected with the FKBP/FRB constructs and treated with rapamycin showed enhanced LRRK2 activity compared to untreated control cells, as measured by a significant increase in the autophosphorylation of LRRK2 at site S1292 as well as phosphorylation of T73 Rab10 and S106 Rab12 (Fig 1B-E). A similar magnitude of LRRK2 activation was also seen with the LYSO-LRRK2 targeting construct (Fig. 1B-E). Thus, directing LRRK2 to lysosomes is sufficient to increase kinase activity, even in the absence of lysosomal damaging agents previously described to activate the LRRK2 pathway (Eguchi et al. 2018; Herbst et al. 2020; Bonet-Ponce et al. 2020).

**Fig 1.**
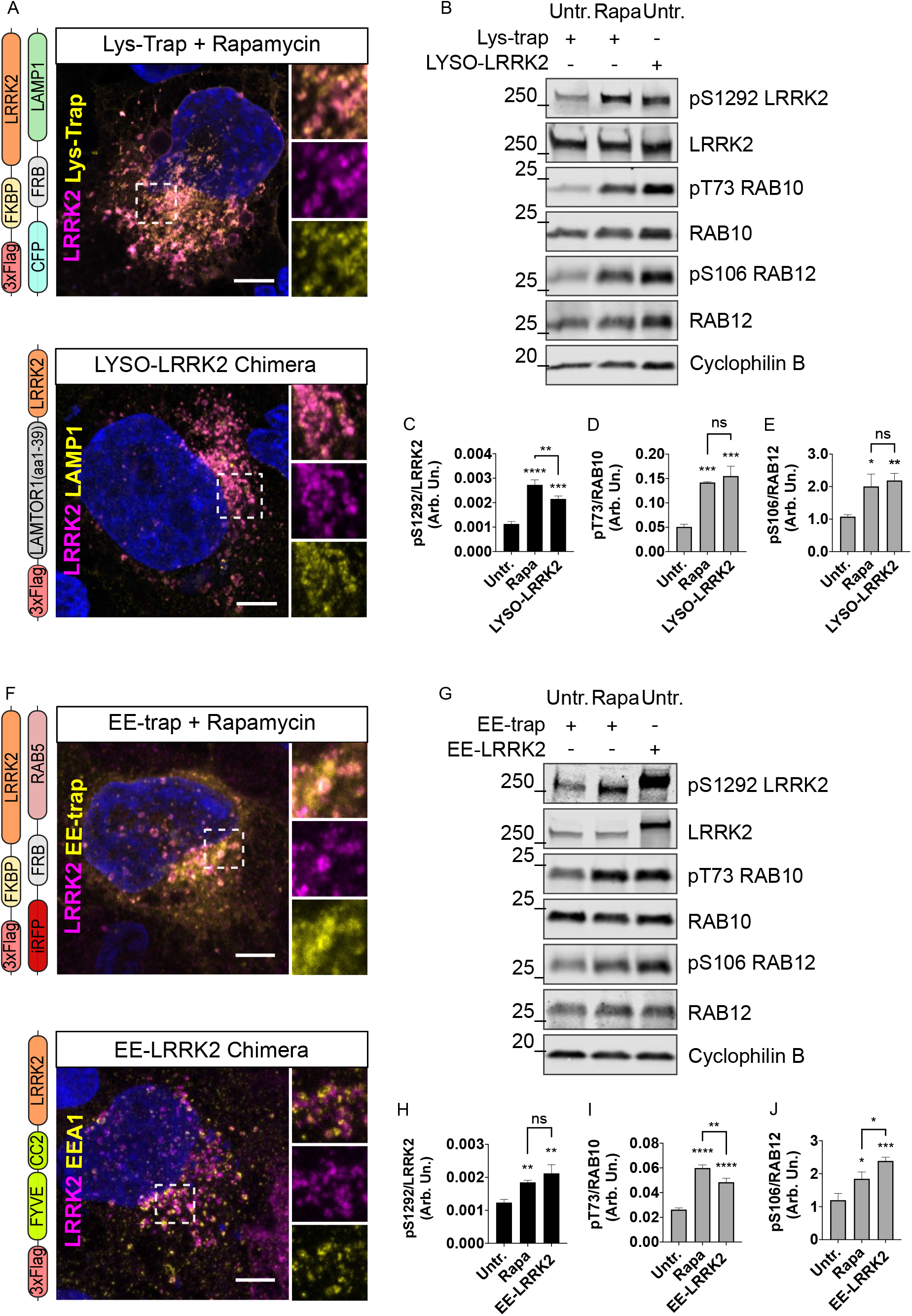
Activation of LRRK2 and subsequent Rab phosphorylation is achieved using FRB-FKBP complex and chimera-LRRK2 constructs regardless of membrane identity. HEK293FT cells transfected with either LAMP1-FRB and FKBP-LRRK2 plasmids (top) or LYSO-LRRK2 plasmid (bottom) (construct designs are shown to the left of the respective immunocytochemistry images in panel A). Scale bar = 10μm. (B-E) Western blot analysis of LRRK2 S1292 autophosphorylation, Rab10 and Rab12 phosphorylation under conditions of Lys-trap with rapamycin treatment or LYSO-LRRK2 chimera transfections. Cells transfected with Rab5-FRB and FKBP-LRRK2 plasmids treated with rapamycin (top) or the EE-LRRK2 chimera (bottom) are shown in panel F (construct designs are shown on the left of the images). Scale bar = 10μm. (G-J) Western blot analysis of LRRK2 S1292 autophosphorylation, Rab10 and Rab12 phosphorylation under conditions of EE-trap with rapamycin treatment or EE-LRRK2 chimera transfections. (C-E, H-J): one-way ANOVA with Tukey’s post hoc; *n* = 3; SD bars shown; (C) *F*(2, 6) = 87.10, (D) *F*(2, 6) = 65.83, (E) *F*(2, 6) = 16.40, (H) *F*(2, 6) = 22.48, (I) *F*(2, 6) = 142.6, (J) *F*(2, 6) = 32.76.

We similarly directed LRRK2 to early endosomal membranes using FKBP-LRRK2 and RAB5-FRB-iRFP constructs and a FYVE-CC2-LRRK2 chimera (referred to as “EE-LRRK2” to reflect an early endosomal chimera). Both methods resulted in LRRK2 colocalization to early endosome markers Rab5 and EEA1, respectively (Fig 1F and Supp. Fig 1D-E). Endogenous VPS35 was also stained as a secondary marker of early endosomes and colocalization with the EE-LRRK2 chimera was quantified (Supplementary Fig 1D-E). Co-transfection of FKBP/FRB-tagged plasmids significantly increased S1292 LRRK2 autophosphorylation and phosphorylation of Rab10 and Rab12 when cells were treated with rapamycin compared to untreated transfected cells (Fig 1G). The EE-LRRK2 chimera similarly activated LRRK2 with results comparable to the rapamycin-treated FKBP/FRB method (Fig 1H-J).

Furthermore, we also targeted LRRK2 to additional membranes using the FKBP/FRB system with FRB-expressed plasmids tagged with markers for late endosomes (LE: iRFP-FRB-RAB7), Golgi (Gol: FRB-CFP-Giantin), recycling endosomes (RE: EHD1-FRB-GFP), and plasma membrane (PM: PM(GAP43_PM-targeting sequence_)-FRB-CFP) in order to interrogate LRRK2 activation across different membrane landscapes (Supp. Fig 2). In the presence of rapamycin, all traps were able to successfully deliver LRRK2 to each specific membrane targeted, determined via colocalization to an endogenous resident protein marker, and to activate LRRK2 kinase as observed by significant increases in pS1292 and downstream pRab10 and pRab12, which were all dephosphorylated when cells were additionally treated with the LRRK2-specific kinase inhibitor MLi-2 (Supp. Fig 2).

Collectively, these results indicate that directing LRRK2 to a given membrane results in activation and phosphorylation of multiple Rab proteins, irrespective of membrane identity. Since both methods of LRRK2 translocation were sufficient for downstream recruitment and phosphorylation of Rabs, we chose to move forward with the lysosome and early endosome chimeric constructs as to avoid unnecessary co-transfections and treatments, i.e. rapamycin, for subsequent experiments.

### LRRK2 and phosphorylated Rab10 colocalization patterns differ at the lysosomal and early endosomal membranes

We next examined the cellular distribution of two LRRK2 substrates, pT73 Rab10 and pS106 Rab12, at endogenous levels using immunostaining. When cells were transfected with LYSO-LRRK2, we noted a significant increase in pT73 Rab10 staining that colocalized to endogenous LAMP2 compared to cells transfected with LRRK2 without lysosomal targeting (referred to as NT-LRRK2 for non-targeted) (Fig 2A). Additionally, when treated with MLi-2, pRab10 signal was completely abolished (Fig 2A-B). Signal intensity of pS106 Rab12 was also significantly increased when LRRK2 is directed to lysosomes via the LYSO-LRRK2 construct compared to NT-LRRK2 and MLi-2 significantly decreased this signal (Fig 2C-D).

**Fig 2.**
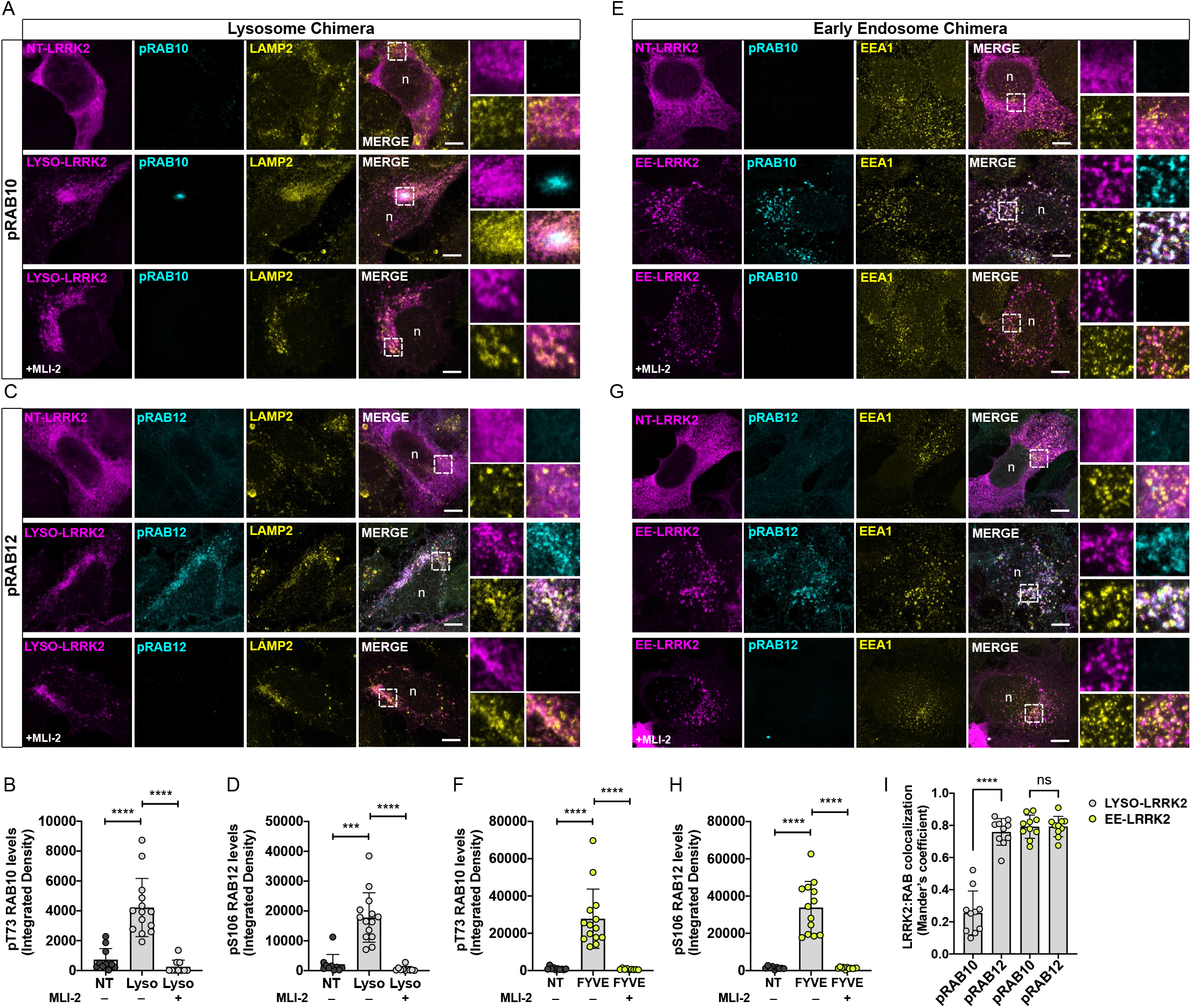
Rab phosphorylation is LRRK2-dependent and pRab colocalization patterns differ between Rab10 and Rab12 at the lysosomal membrane. HEK293FT cells were transfected with either NT-LRRK2 or LYSO-LRRK2 chimera (magenta) and pT73 Rab10 and pS106 Rab12 (cyan) integrative densities were measured (A-D). MLi-2 treatment was compared to untreated cells expressing LYSO-LRRK2 (A-D). Scale bar = 10μm. (E-I) Cells were transfected with either NT-LRRK2 or EE-LRRK2 chimera (magenta) and pT73 Rab10 and pS106 Rab12 (cyan) integrative densities were measured (A-D). MLi-2 treatment was compared to untreated cells expressing EE-LRRK2 (E-I). Scale bar = 10μm. (B, D, F, H): one-way ANOVA with Tukey’s post hoc; (I): two-way ANOVA with Tukey’s post hoc; N=2, *n* = 13 cells; SD bars shown; (B) *F*(2, 32) = 33.11, (D) *F*(2, 32) = 34.92, (F) *F*(2, 32) = 26.67, (H) *F*(2, 32) = 52.36, (I) Chimera: *F*(1, 36) = 93.49, *p*<0.0001, Rab: *F*(1, 36) = 73.37, *p*<0.0001.

When cells were transfected with the EE-LRRK2 construct, both phosphorylation of Rab10 and Rab12 were significantly increased and colocalized with endogenous EEA1 compared to the transfection of the untagged LRRK2 construct, and the addition of MLi-2 caused a significant reduction in signals for both pRabs (Fig 2E-H). These data suggest that Rab10 and Rab12 are phosphorylated by LRRK2 and can be recruited to any membrane where LRRK2 kinase is present, likely due to a lack of removal by GDI (Rab GDP dissociation inhibitor) as previously suggested (Gomez et al., 2019).

Inspection of the same images suggested that not all LRRK2-positive lysosomes were also pRab10 positive (Fig. 2A). Therefore, we next measured LRRK2:pRab colocalization at LRRK2-positive lysosomes and LRRK2-positive early endosomes using Mander’s coefficient. Quantitatively, we found that only a small portion of LRRK2-positive lysosomes colocalized to pT73 Rab10, whereas the majority of LRRK2-positive lysosomes showed colocalization to pS106 Rab12 (Fig 2I). Additionally, nearly all LRRK2-positive early endosomes colocalized with both pRab10 and pRab12 (Fig 2I). We also observed that this cluster of pRab10 lysosomes was situated in the perinuclear region, and not observed distally. Staining for total Rab10 on cells transfected with LYSO-LRRK2 revealed a similar pattern to pRab10, in which the total protein colocalized only to a subset of perinuclear LRRK2-positive lysosomes (Supp. Fig 3A-B). Additionally, we stained for JIP4, a motor adaptor protein recruited by phosphorylated Rab10 to the lysosomal membrane (Waschbüsch et al. 2020; Bonet-Ponce et al. 2020) and found it to also cluster with the subset of LRRK2-positive lysosomes on cells expressing LYSO-LRRK2 compared to NT-LRRK2 (Supp. Fig 3C-D). This data suggests that while Rab10 is retained at LRRK2-positive lysosomes after phosphorylation, there are additional conditions that determine why pRab10 is present at a subset of lysosomes in the perinuclear region of the cell.

### Expression of motor proteins ARL8B and SKIP promote dispersion of LRRK2-positive lysosomes to the cell periphery

In order to try to delineate the mechanism underlying the recruitment and phosphorylation of Rab10 to a subset of lysosomes, we took a spatial approach by manipulating lysosomal position. The small Arf-like GTPase ARL8B and kinesin-1 adaptor protein SKIP have been shown to direct lysosomes to the cell periphery when overexpressed (Rosa-Ferreira and Munro 2011; Keren-Kaplan and Bonifacino 2021). Briefly, ARL8B binds to both the lysosomal membrane and SKIP, which contains two light chain kinesin-binding domains to promote anterograde movement along microtubules (Fig 3A). Interestingly, we found considerable morphological changes to HEK293FT cells with the addition of the ARL8B and SKIP proteins in which cells formed long processes with a large accumulation of lysosomes situated at the tips (Fig 3B-C). When measuring the number of LRRK2-positive lysosomes at the periphery versus total lysosome count, a significant increase was observed in the number of LRRK2-positive lysosomes situated at the periphery when expressing ARL8B and SKIP together (Fig 3D).

**Fig 3.**
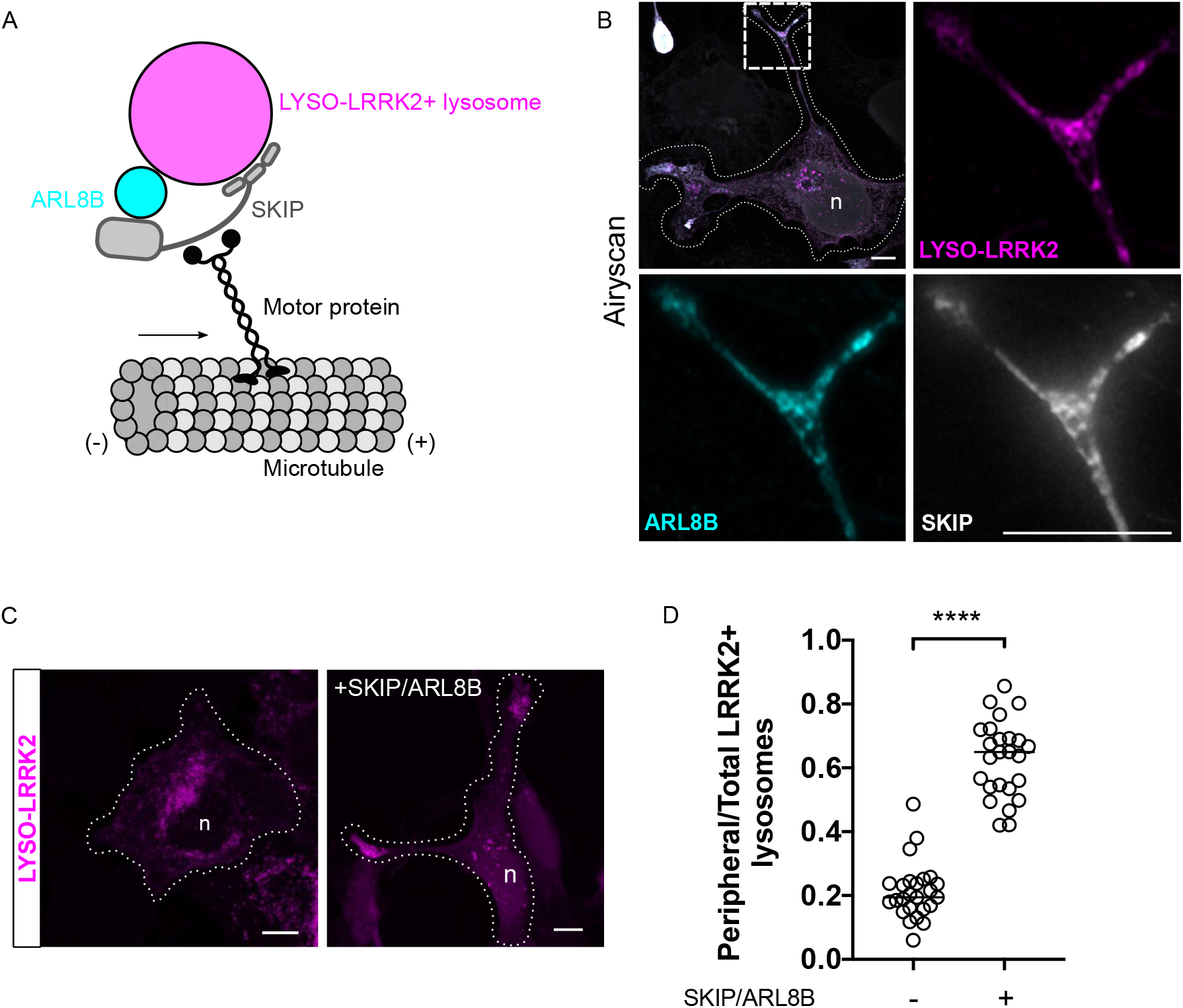
ARL8B- and SKIP-expressing cells promote lysosomal repositioning to the periphery of cells. Representative schematic describing lysosomal translocation to the cell periphery via expression of ARL8B and SKIP proteins along the plus end of microtubules (A). Airyscan microscopy showing peripheral LYSO-LRRK2 lysosomes (magenta), colocalizing with ARL8B (cyan) and SKIP (grey) (B). Cell edges are outlined. Scale bar of whole cell= 20μm; scale bar of boxed close up= 2μm. LYSO-LRRK2 (magenta) was transfected in cells alone or co-transfected with ARL8B and SKIP (C). Scale bar= 10μm, 15μm. Quantification of the ratio of peripheral to total LRRK2-positive lysosomes such as those represented in panel C are shown in (D). Peripheral fluorescence refers to the presence of LRRK2 within 2 μm from the cell vertices. Horizontal lines indicate the mean SD from 3 independent experiments.

### Phosphorylated Rab10 does not colocalize to peripheral LRRK2-positive lysosomes while lysosomal clustering towards the perinuclear area increases colocalization

We next co-transfected LYSO-LRRK2, ARL8B, and SKIP and stained for pT73 Rab10 to see whether lysosomes situated at the periphery would be able to phosphorylate and retain Rab10. Both pRab10 levels and colocalization with LRRK2 were significantly decreased when cells were transfected with ARL8B and SKIP compared to cells transfected with LYSO-LRRK2 alone (Fig 4A-C). This pattern was also observed when staining for total Rab10 levels (Supp. Fig 3E). Taken together, these data show that lysosomes situated within the perinuclear region have specific properties that allow recruitment and phosphorylation of Rab10 by LRRK2 that are not shared with peripheral lysosomes.

**Fig 4.**
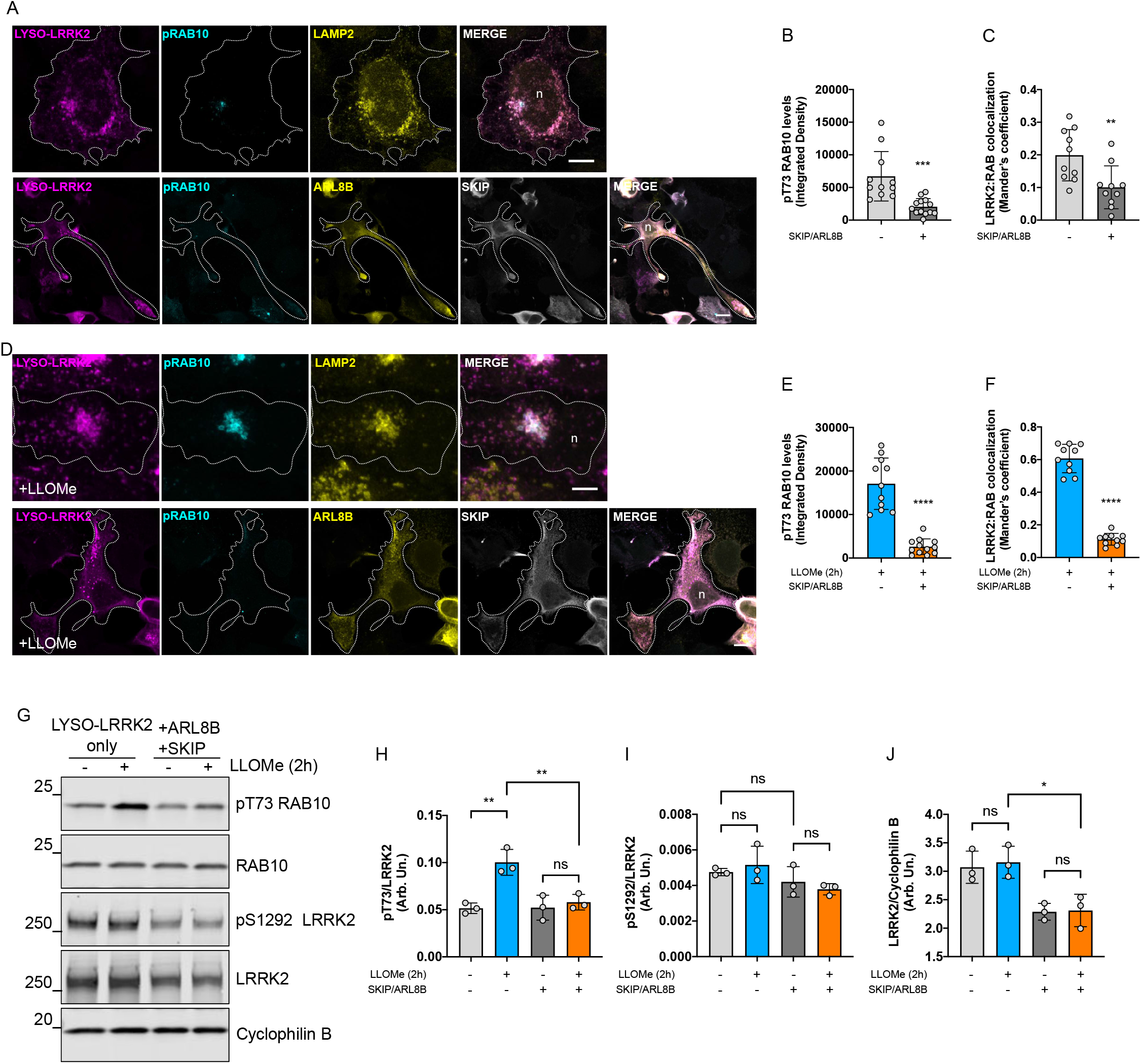
ARL8B- and SKIP-expressing cells prevent pRab10 colocalization on peripherally positioned LRRK2-positive lysosomes. Representative confocal microscopy images show pT73 Rab10 signal (cyan), LYSO-LRRK2 (magenta) with endogenous LAMP2 (yellow) (top) or ARL8B (yellow) and SKIP (grey) (bottom) (A). Integrative density and Mander’s correlation coefficient were used to measure pRab10 staining intensity (B) and LRRK2:pRab10 colocalization (C). (D) Cells with and without ARL8B/SKIP transfection were treated with 1 mM LLOMe for 2 hours. Integrative density and Mander’s correlation coefficient were used to measure pRab10 staining intensity (E) and LRRK2:pRab10 colocalization (F). LYSO-LRRK2 expressing cells with and without ARL8B/SKIP co-transfection and LLOMe treatment were probed for S1292 LRRK2 and pT73 Rab10 for Western blot analysis (G-J). (B-C, E-F): two-tailed unpaired t test; n= 13 cells across two independent experiments; error bars represent SD; (B) \*\*\**p*= 0.0004, (C) \*\**p*=0.0071, (E) \*\*\*\**p*<0.0001, (F) \*\*\*\**p*<0.0001. (H-J): one-way ANOVA with Tukey’s post hoc; *n*= 3; SD bars shown. (H) *F*(3,8) = 14.17, (I) *F*(3,8) = 2.201,(J) *F*(3,8) = 10.16. Scale bars = 10 μm, 5 μm.

Previously, we have shown that the addition of a lysosomotropic agent LLOMe induces LRRK2 localization to the lysosomal membrane and recruitment of Rab10 in order to initiate a lysosomal tubulation process (LYTL) via JIP4 (Bonet-Ponce et al. 2020). We thus conjectured that adding LLOMe to cells transfected with SKIP and ARL8B would push Rab10 to LRRK2-positive lysosomes despite their location at the periphery. However, acute treatment with LLOMe did not affect the recruitment nor subsequent phosphorylation of Rab10 in cells transfected with SKIP and ARL8B (Fig 4D-F). Western blot analysis supported these results further, in that LLOMe treatment significantly increases phosphorylation of Rab10 in cells transfected only with LYSO-LRRK2 but this effect was counteracted by co-transfection with ARL8B and SKIP (Fig 4G-H). Interestingly, we do not observe any impact of LRRK2 autophosphorylation at S1292 upon treatment with LLOMe (Fig 4G, I), suggesting that LLOMe induces LRRK2 in a manner that is distinct from LRRK2 mutations that all increase pS1292.

When staining for pRab12 in cells transiently expressing ARL8B and SKIP, we found strong colocalization between pRab12 and LRRK2-positive peripheral lysosomes regardless of lysosomal position, and acute treatment with LLOMe equally increased pRab12 in cells transfected with LYSO-LRRK2 alone or with ARL8B and SKIP (Fig 5A-D). This suggests that there are differing mechanisms at play for LRRK2-dependent phosphorylation of Rab10 and Rab12 at lysosomes. Interestingly, we also measured pT71 Rab29 levels for conditions under LLOMe treatment at peripheral and perinuclear lysosomes and found a similar pattern to that of pRab10, in which peripherally sequestered lysosomes are unable to phosphorylate Rab29 (Fig 5E-F).

**Fig 5.**
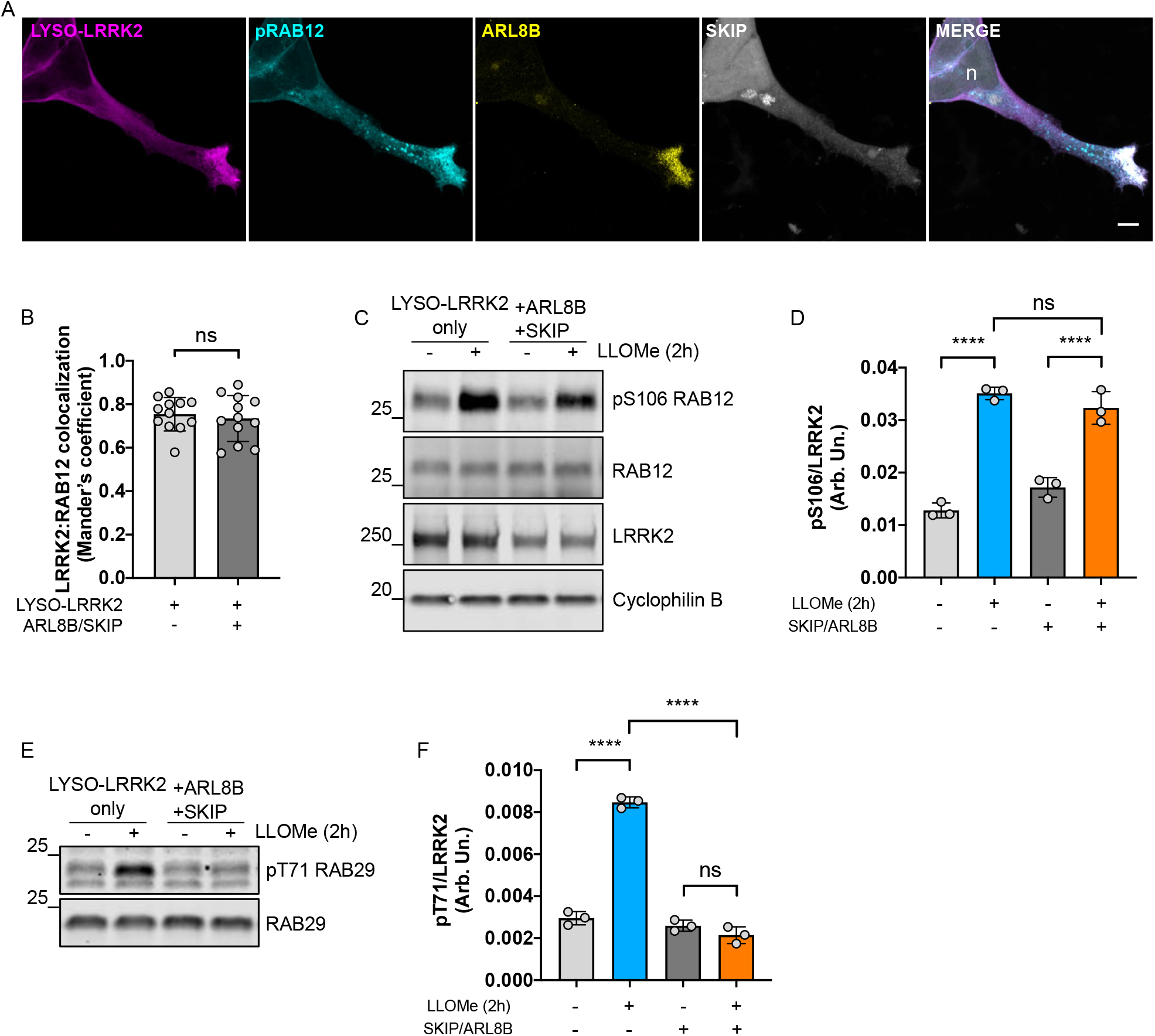
Phosphorylated Rab12 is found at peripheral lysosomes in cells transiently transfected with ARL8B and SKIP plasmids. A super-resolution confocal microscopy image shows pS106 Rab12 staining (cyan) at LRRK2-positive lysosomes (magenta) when ARL8B (yellow) and SKIP (grey) are co-expressed (A). Scale bar= 20μm. Mander’s coefficient was used to measure LRRK2:pRab12 colocalization in conditions where ARL8B and SKIP are co-expressed with LYSO-LRRK2 as well as when only LYSO-LRRK2 is expressed alone (B). Western blot analyses of Rab12 and Rab29 phosphorylation are shown (C-F). (B): Unpaired, two-tailed t test; N=2, n= 12 cells each; SD error bars are shown; *p*= 0.6045. (D-F): one-way ANOVA with Tukey’s post hoc; *n*=3; SD bars shown. (D) *F*(3,8) = 87.73, (F) *F*(3,8) = 268.0.

To further uncover the mechanism behind position-specific Rab10 phosphorylation on LRRK2-positive lysosomes, we turned our attention to JIP4. Although we have previously shown that JIP4 binds to pRab10 to induce lysosomal tubulation, JIP4 is a motor adaptor protein that binds to the dynein/dynactin complex to promote transport of lysosomes towards the minus end of microtubules (Willett et al. 2017) (Fig 6A). Using siRNA, we knocked down JIP4 followed by transient transfection of LYSO-LRRK2 and stained for pRab10. We confirmed similar lysosomal dispersion to the periphery as seen with ARL8B and SKIP overexpression and saw a significant decrease in pRab10 signal compared to the non-targeting control siRNA condition (Fig 6B-C). This finding identifies that JIP4 influences lysosomal positioning and thus LRRK2-dependent phosphorylation of Rab10.

**Fig 6.**
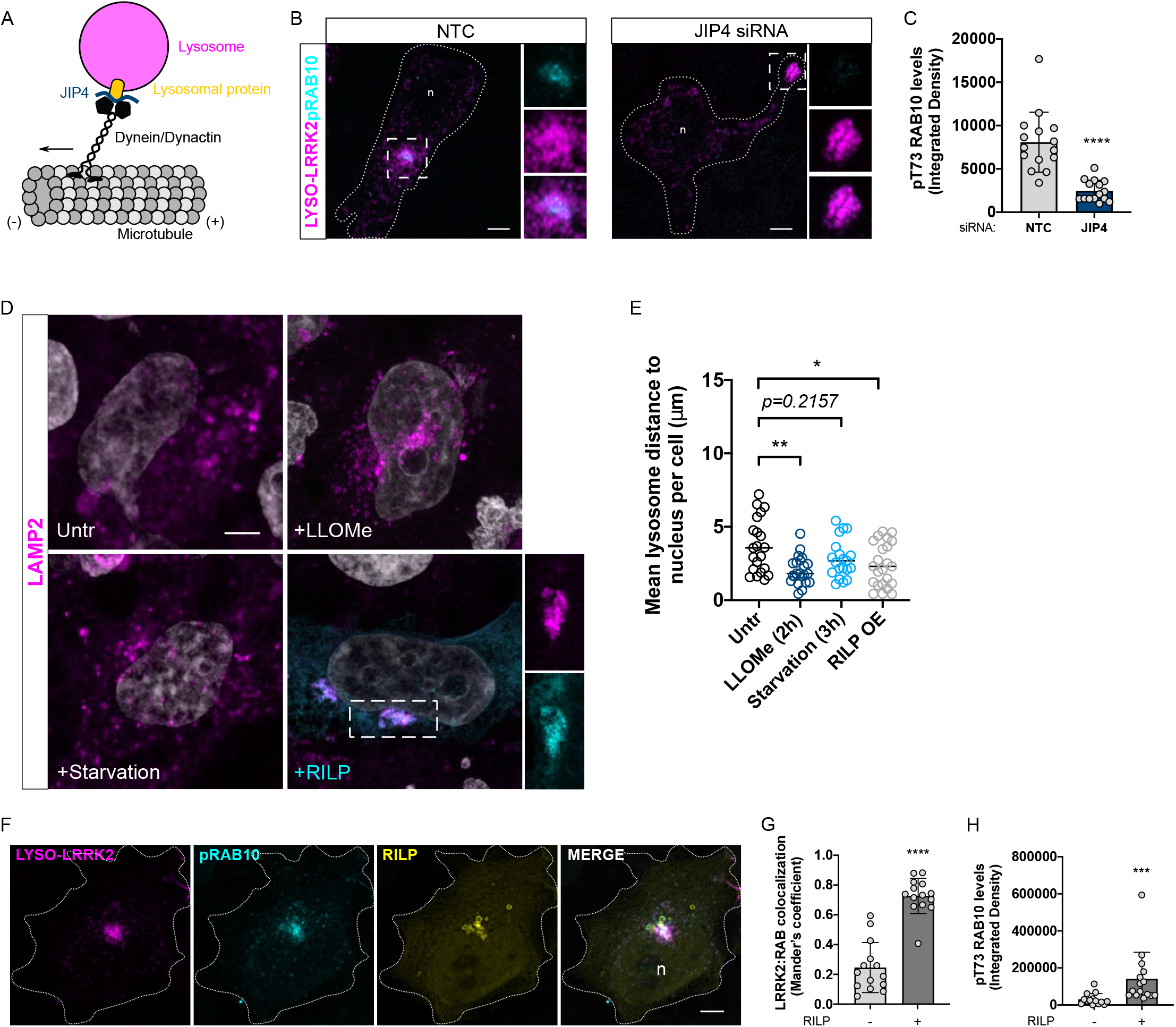
RILP-expressing cells promote clustering of LRRK2-positive lysosomes within the perinuclear region and significantly increase pRab10 signal. Simplified schematic of lysosomal movement into the perinuclear area via JIP4 adaptor binding to a lysosomal protein and the dynein/dynactin complex (A). Representative confocal microscopy images of HEK293FT cells stained for LYSO-LRRK2 (magenta) and pRab10 (cyan) under conditions of NTC and JIP4 siRNA transfection (B), in which knockdown of JIP4 significantly reduces pRab10 signal (C). Representative confocal microscopy images for a set of experiments showing methods of lysosomal clustering in HEK293FT cells (A). Cells were subjected to conditions of 2 hours LLOMe treatment, 3 hours serum and amino acid starvation in EBSS, or overexpression of RILP. Endogenous LAMP2-positive lysosomes (magenta) were counted and the shortest distance to the nucleus was measured per lysosome (D). These values were then averaged per cell (E). Cells expressing LYSO-LRRK2 (magenta) and RILP (yellow) were stained for pRab10 (cyan) (F). Nearly 80% of LRRK2-positive lysosomes were colocalized to pRab10 staining with RILP overexpression compared to control cells (G). Integrated density of pRab10 signal was significantly increased in cells expressing RILP compared to control cells without RILP (H). (C, G-H): two-tailed unpaired t test; n= 15 cells. SD error bars are shown; (C) \*\*\*\**p*<0.0001, (G) \*\*\*\**p*<0.0001, (H)***p= 0.0003. (E): one-way ANOVA with Tukey’s post hoc; n= 25 cells across 3 independent experiments; horizontal line represents the mean SD. (E) *F*(3,83) = 4.642.

LLOMe treatment not only works as a lysosomal membrane damaging reagent, but also promotes the clustering of lysosomes in the perinuclear region and increasing Rab10 phosphorylation in the perinuclear region as a consequence. Therefore, we compared different stimuli to promote lysosomal retrograde transport in HEK293FT cells, including LLOMe treatment, RILP overexpression, and serum/amino acid starvation. RILP is a Rab7A effector that affects lysosomal positioning within the cell via recruitment of dynein and dynactin proteins (Cantalupo et al. 2001). Overexpressing RILP arrests Rab7A in its GTP-bound active form and sequesters the protein to late endosomal and lysosomal membranes and induces recruitment of the dynein-dynactin complex which promotes vesicle translocation along the minus end of microtubules towards the perinuclear region (Jordens et al. 2001). We found that both LLOME treatment and RILP overexpression significantly increased the clustering of lysosomes into the perinuclear area, whereas starvation did not have a significant effect on lysosomal clustering as determined by measuring the mean distance of endogenous LAMP2 puncta to the nucleus in HEK293FT cells (Fig 6D-E).

Based on these results, we co-expressed LYSO-LRRK2 and RILP and stained for pT73 Rab10. Excitingly, we found that RILP overexpression is sufficient to cluster LRRK2-positive lysosomes into the perinuclear region, resulting in the recruitment and phosphorylation of Rab10, as demonstrated by LRRK2:pRab10 colocalization (Fig 6F-G). We also observed phosphorylation levels of Rab10 were significantly increased compared to cells that were transfected with LYSO-LRRK2 alone (Fig 6H). These results further demonstrate that lysosomes situated at the perinuclear area possess favorable characteristics for the recruitment and phosphorylation of Rab10 by LRRK2. Additionally, we stained for pericentrin, a marker for centrosomes, to determine where the pRab10-positive lysosomes were in relation to the centrosome. We found that these lysosomes were situated adjacent to pericentrin, but did not colocalize with pericentrin in both HEK293FT and U2OS cell types (Supp. Fig 4A-B).

### Dispersion of lysosomes to the cell periphery is sufficient to block recruitment of Rab10 in untagged LRRK2 and endogenous LRRK2 models

We next wanted to test whether the phenomenon of peripheral lysosomes being recalcitrant to pRab10 formation could be recapitulated using LLOMe treatment in non-targeted, wildtype LRRK2 expressing cells. NT-LRRK2 is primarily cytosolic, and treatment with LLOMe promotes colocalization to the lysosomal membrane (Herbst et al. 2020; Bonet-Ponce et al. 2020). Therefore, we treated cells transfected with NT-LRRK2 with LLOMe. We observed that LRRK2 relocalized to the lysosomal membrane with or without the addition of ARL8B and SKIP overexpression, shown as colocalization with endogenous LAMP2 (Fig 7A-B). Interestingly, when cells were treated with LLOMe, colocalization of LRRK2 and pRab10 was found only in the subset of lysosomes within the perinuclear region and this was blocked when lysosomes are pushed to the periphery upon ARL8B and SKIP overexpression (Fig 7C-D). Furthermore, JIP4 recruitment to LLOMe-treated lysosomes was also prevented by ARL8B- and SKIP expression (Fig 7E-F), suggesting that LYTL can not be activated at peripheral lysosomes. Western blot analysis confirmed that the addition of SKIP and ARL8B inhibits pRab10 in cells treated with LLOMe compared to those without ARL8B and SKIP overexpression (Fig 7G-I). Interestingly, the movement of most lysosomes to the periphery does not affect the ability of Galectin-3 recruitment to damaged lysosomes after treatment with LLOMe, suggesting that lysophagy is not inhibited at the periphery (Supp. Fig 5).

**Fig 7.**
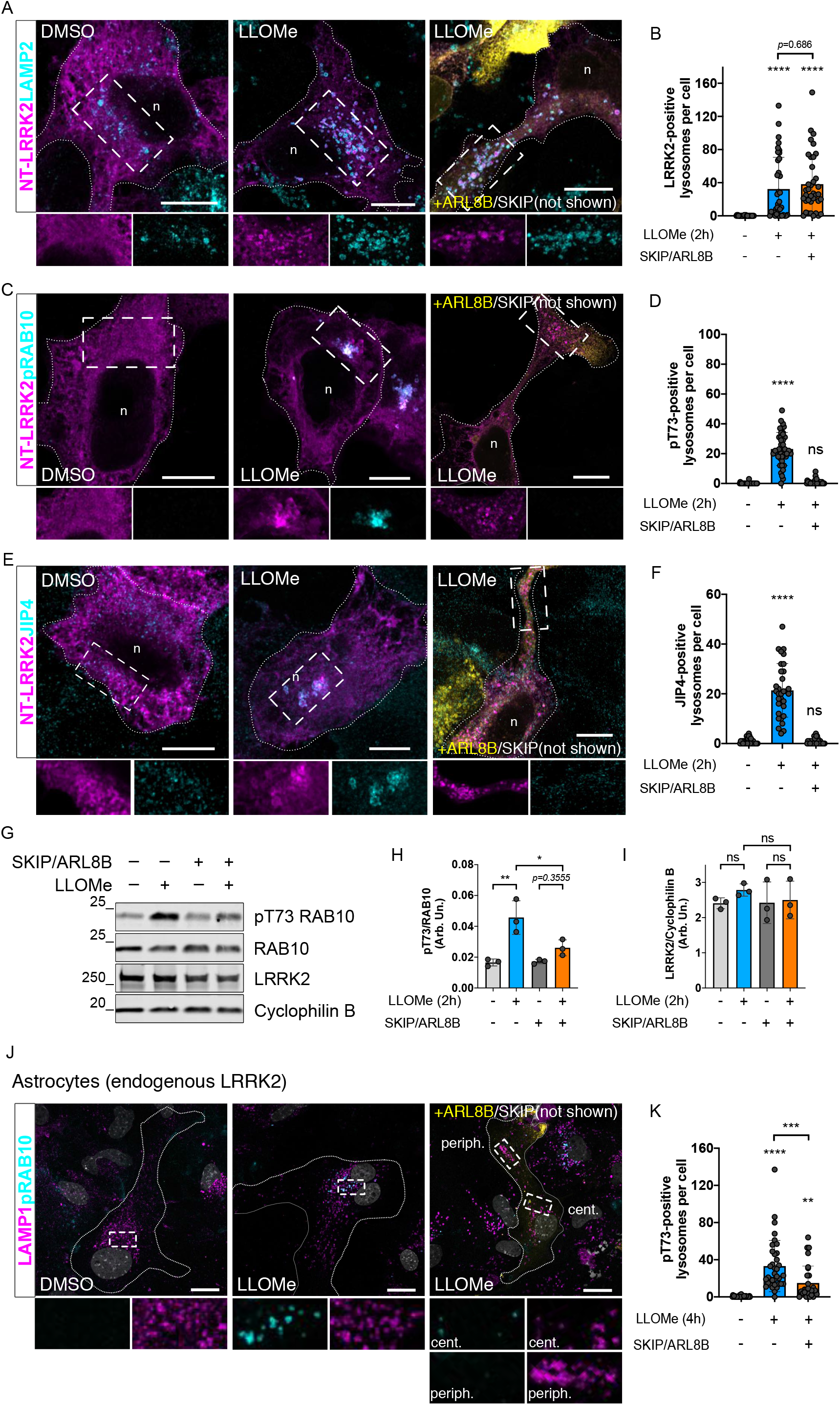
ARL8B- and SKIP-expressing cells treated with LLOMe recruit NT-LRRK2-expressing HEK293FT and endogenous LRRK2 in primary mouse astrocytes but not pRab10. Representative Airyscan microscopy images of NT-LRRK2 (magenta) with and without ARL8B/SKIP (yellow) co-expressing cells under conditions of DMSO or LLOMe treatment. (A-B) LRRK2-positive lysosomes are counted in each condition (LAMP2 endogenous marker in cyan), followed by pT73 Rab10 (C-D), and lastly JIP4 (E-F). Scale bars= 8μm, 10 μm, 15 μm. (G-I) Western blot analysis of pRab10 from whole sample lysates of LYSO-LRRK2 expressing cells +/− ARL8B/SKIP and +/− LLOMe. (J-K) Primary astrocytes treated with LLOMe resulted in a strong increase in pRab10 signal compared to DMSO-only treated cells. This signal was found to be lysosome-positive (as determined via endogenous LAMP1 colocalization) (J-K). Of the remaining pRab10 signal, all was found within the perinuclear region compared to lysosomes that were peripherally situated (last image in panel J). Scale bar= 20μm. (B,D,F,H-I,K): one-way ANOVA with Tukey’s post hoc; (B,D,F,K) n = 32-40 cells counted. (H-I) n= 3; SD error bars are shown. (B) *F*(2,105) = 15.91, (D) *F*(2,107) = 141.1, (F) *F*(2,94) = 116.4, (H) *F*(3,8) = 14.54, (I) *F*(3,8) = 0.5329, (K) *F*(2,99) = 27.14.

We then decided to evaluate whether the same effects were seen with LRRK2 at endogenous levels, and selected mouse primary astrocytes for their high levels of LRRK2 expression (Bonet-Ponce et al. 2020; Alexandra Beilina et al. 2020). Acute LLOMe treatment for 4 hours significantly increased phosphorylated Rab10 at lysosomes compared to those only treated with DMSO (Fig 7J-K), whereas ARL8B and SKIP overexpression resulted in a significant reduction in pRab10-positive lysosomes, and minimal recruitment to peripheral lysosomes was observed (Fig 7J-K). Taken together, our data indicate that LRRK2-dependent Rab10 recruitment to the lysosomal membrane is position-specific, and is recapitulated across multiple models including at endogenous levels of LRRK2.

### Knockdown of PPM1H increases phosphorylation of Rab10 at perinuclear lysosomes and increases the frequency of lysosomal tubulation via JIP4

A recent siRNA screen identified that PPM1H phosphatase dephosphorylates T72 Rab8A and T73 Rab10, thus mitigating LRRK2 signaling (Berndsen et al. 2019). Of interest, exogenously expressed PPM1H is reported to be located at the Golgi, thus potentially having spatially restricted effects on cellular phosphorylation. We therefore hypothesized that PPM1H might counteract LRRK2-dependent Rab10 phosphorylation on lysosomes in the perinuclear area. Knockdown of PPM1H resulted in a significant increase in pRab10 signal on LYSO-LRRK2 lysosomes within the perinuclear region but no increase in signal at peripheral lysosomes was observed (Fig 8A-B). A similar pattern was seen when quantifying JIP4 signal as expected (Fig 8C-D). Under these conditions, we also found a significant increase in the frequency of pRab10- and JIP4-positive lysosomal tubules (Fig 8E-H). This finding demonstrates that PPM1H limits pRab10 phosphorylation at the perinuclear lysosomes and thereby counteracts LYTL.

**Fig 8.**
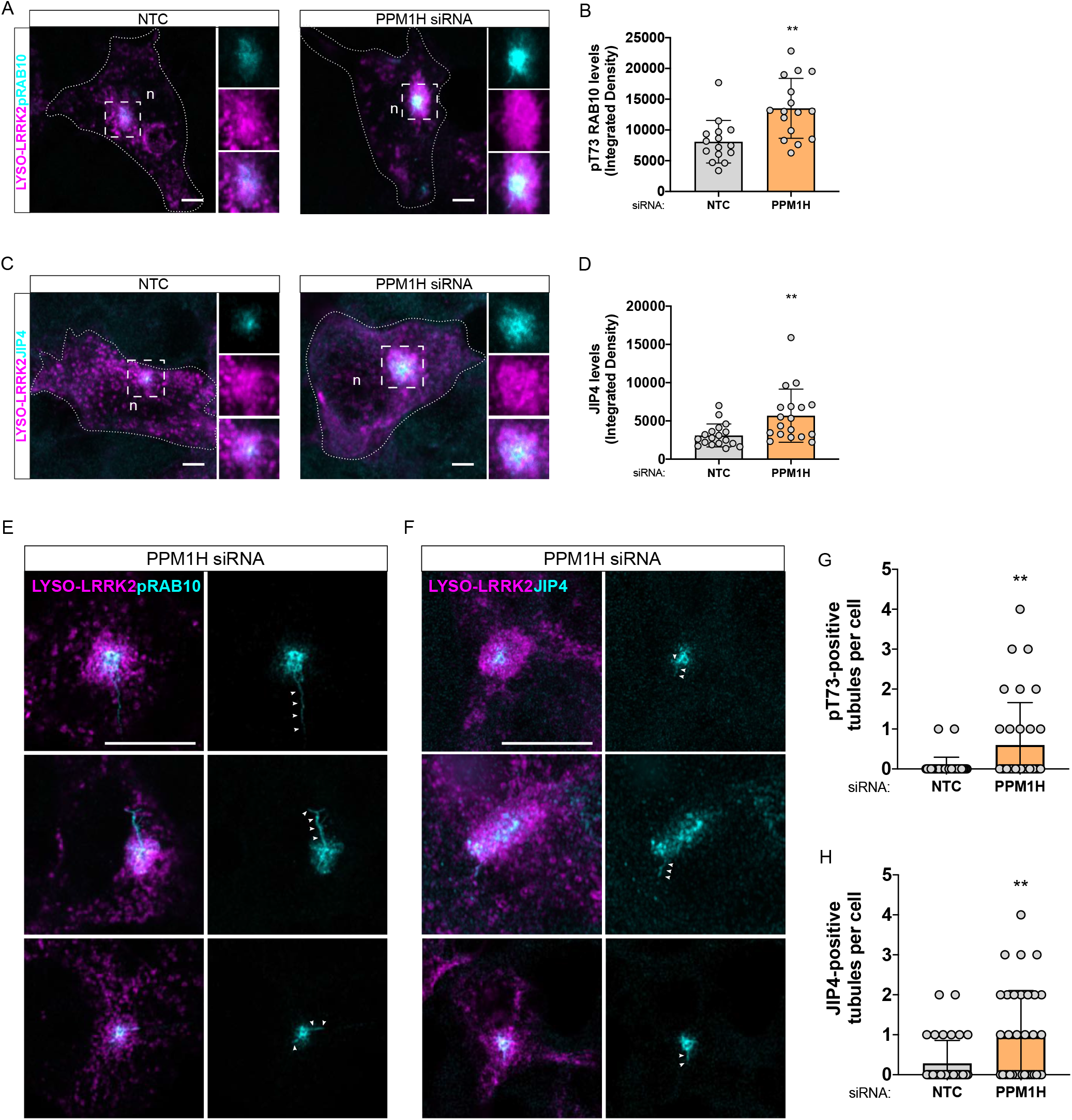
PPM1H knockdown affects pRab10-mediated lysosomal tubulation. Representative confocal images showing LYSO-LRRK2 (magenta) and pT73 Rab10 (cyan) or JIP4 after siRNA knockdown of NTC or PPM1H (A, C). Airyscan images are shown of pT73 Rab10- and JIP4-positive tubules (E-F). (B, D, G-H): two-tailed unpaired t test; n=16 (B, D); n=35 (G-H); SD bars shown. (B) p=0.0013, (D) p=0.0066, (G) p=0.0055, (H) p=0.0038.

## DISCUSSION

Understanding the patterns of LRRK2 kinase activity is of crucial interest for PD pathogenesis and future therapeutics targeted at this enzyme. Recent work from our lab and others have shown that LRRK2 can be targeted to different membranous compartments where it can phosphorylate and recruit several Rab substrates under specific cell signaling conditions. For example, LRRK2 has been robustly shown to translocate to the TGN, the phagosomal membrane and the membrane of damaged lysosomes depending on the stimulus used (Beilina et al. 2014; Herbst et al. 2020; Eguchi et al. 2018; Purlyte et al. 2019; Bonet-Ponce et al. 2020; Lee et al. 2020). However, the dynamics underlying LRRK2 membrane presence, activation and Rab phosphorylation remain unclear and, specifically, whether membrane association is necessary or sufficient for LRRK2 activation. In the current study, we have shown that LRRK2 dependent cellular pathways depend on multiple regulatory events and that downstream Rab phosphorylation is influenced by lysosomal positioning and the action of spatially restricted phosphatases.

We found that LRRK2 undergoes activation, as measured by autophosphorylation at S1292, irrespective of the membranous compartment. Additionally, we noted a similar increase in phosphorylation of T73 Rab10 and S106 Rab12 at all membranes studied, suggesting first, that membrane identity is not important for LRRK2 activation and, second, that pRabs do not seem to exhibit membrane preferences, but can be retained at any membrane once phosphorylated by LRRK2. This data is consistent with recent observations using a mitochondrial Rab29 chimera to drive LRRK2 to the mitochondrial membrane that was suggested to occur due to GDI1/2 proteins being unable to remove phosphorylated Rabs away from membranes (Gomez et al. 2019).

Additionally, we characterized pRab localization patterns after LRRK2 was translocated to the lysosomal and early endosomal membranes. We observed a striking difference in colocalization pattern between Rabs, with pT73 Rab10 showing strict perinuclear localization in a subset of lysosomes whereas pS106 Rab12 was found in LRRK2-positive lysosomes throughout the cell. Recently, we proposed that pRab12 more accurately reports the effects of inhibition of LRRK2 kinase activity than pRab10 *in vivo* (Kluss et al. 2021). Our current data suggests that Rab12 is regulated by additional mechanisms distinct from Rab10 (discussed below). Interestingly, JIP4 showed a perinuclear localization that was similar to pRab10 in cells treated with LLOMe, confirming that the induction of LYTL occurs after pRab10 accumulates on juxtanuclear lysosomes.

We have identified a spatially-dependent, lysosomal-specific mechanism through which LRRK2 recruits and phosphorylates Rab10. When LRRK2-positive lysosomes are pushed to the periphery via ARL8B and SKIP overexpression, Rab10 can no longer be recruited, in contrast to Rab12. In contrast, pRab10 colocalization to LRRK2 was significantly increased when lysosomes were clustered into the perinuclear region using LLOMe treatment or RILP overexpression.This phenomenon was also seen with knockdown of endogenous JIP4, revealing a novel role of JIP4 in LRRK2-dependent phosphorylation of Rab10 at lysosomes positioned near the nucleus. We speculate that there may be a possible feedforward signaling pathway where lysosomal positioning can be promoted by JIP4 to the perinuclear region that would then localize JIP4 for initiation of LYTL after phosphorylation of Rab10.

What remains to be identified are the key conditions that determine spatial-specificity of phosphorylation of specific Rabs at lysosomes. Lysosomes are highly dynamic and transient organelles responsible for wide ranging cellular functions such as autophagy, nutrient response, cell growth and migration (Cabukusta and Neefjes 2018), and clustering lysosomes into the organelle-dense perinuclear cloud has been observed under various conditions of cellular stress and compromisation (Korolchuk et al. 2011; de Martín Garrido and Aylett 2020). Cells are thought to relocalize lysosomes to provide optimal conditions for handling membrane damage, including transfer of undegraded materials to newly synthesized lysosomes, as well as to promote catalytic activity and induce lysophagy of damaged lysosomes. LYTL, exacerbated by LLOMe treatment, requires the recruitment and phosphorylation of Rab10 and its subsequent recruitment of JIP4 to induce tubulation, which we previously hypothesized was beneficial to sort undegraded cargo to other active lysosomes for proper disposal (Bonet-Ponce et al. 2020). Our data collectively suggest that at least two conditions are required for pRab10/JIP4 presence on lysosomes: (1) LRRK2 must be active and recruited to lysosomes, and (2) LRRK2-positive lysosomes must be located near the nucleus. Thus, the current data suggest a complex regulation of signaling downstream of LRRK2 that involves multiple conditions and varies depending on which Rab is phosphorylated by LRRK2.

Recent work by Berndsen et al. identified T73 Rab10 as a substrate for PPM1H phosphatase, thus counteracting LRRK2-signaling (Berndsen et al. 2019). Here, we show that knockdown of PPM1H significantly increased pRab10 signal as well as pRab10- and JIP4-positive tubules on perinuclear lysosomes, while LRRK2-positive peripheral lysosomes were unaffected. PPM1H has been suggested to be resident to the Golgi and therefore may only influence Rabs within the perinuclear space. By extension, this suggests that as of yet unidentified phosphatases may influence Rab10 phosphorylation at peripheral lysosomes and we infer that these phosphatases are likely to be highly active. Of note, Berndsen et al. reported that pS106 Rab12 was not influenced by PPM1H. If different phosphatases act on selected pRabs, we speculate that the dissociation between the effects of lysosomal positioning between Rab10 and Rab12 arise from the differential actions of these, again unidentified, phosphatases.

Overall, our results suggest that the presence of LRRK2 to a membrane is sufficient to trigger its activation, Rab phosphorylation and retention at membranes, and that Rab10-specific recruitment to lysosomes is, in part, controlled by lysosomal positioning. Further studies are needed in order to identify the key proteins involved in determining optimal conditions for LRRK2-dependent Rab10 recruitment at the lysosomal membrane, including the description of novel phosphatases that can act on Rab12 and on Rab10 at peripheral lysosomes. Our results suggest LRRK2 signaling pathways are complex and will function as a product of which Rab is phosphorylated and which phosphatases act to counteract or terminate that signal.

## MATERIALS AND METHODS

### Cell culture

HEK293FT cells and primary astrocytes were maintained in DMEM, or DMEM/F12, respectively, containing 4.5 g/l glucose, 2 mM l-glutamine, 5% Pen/Strep and 10% FBS at 37°C in 5% CO_2_. For experiments, cells were seeded on 12 mm coverslips pre-coated with matrigel. Primary astrocytes were cultured for 2 weeks from P0 pups and split twice before usage.

### Reagents and treatments

For all experiments utilizing the FKBP-FRB complex, rapamycin (Cayman Chemicals, cat #13346) was added at 200 nM for 15 min prior to fixation in 4% PFA. MLi-2 (Tocris, cat #5756) was used at 1 μM in DMSO for 90 min, or LLOMe (Sigma-Aldrich, cat #L7393) was added at 1 mM in DMSO prior to fixation or lysing cells for downstream analyses. Starvation experiments consisted of rinsing cells with PBS followed by incubation in EBSS for 3 hours before fixation. Time points from 2 hours to 16 hours of starvation were explored and all resulted in no significant lysosomal clustering in HEK293FT cells.

### Cloning

The FKBP sequence was tagged to a 3xFLAG-LRRK2 vector using IN-FUSION HD cloning technology (Clontech, Takara, cat #638920). LYSO-LRRK2 was created by adding the N terminal domain of the LAMTOR1 sequence (aa 1-39) into the 3xFLAG-LRRK2 plasmid using IN-FUSION HD cloning technology (LAMTOR1(1-39)-3xFLAG-LRRK2). FYVE-LRRK2 was created by cloning the FYVE-CC2 targeting sequence of HRS into a 3xFLAG pDEST using IN-FUSION HD cloning technology. NT-LRRK2, previously cloned into a pCR™8/GW/TOPO™ vector (ThermoFisher, cat #250020), was then transferred into the new 3xFLAG-FYVE-CC2-pDEST using Gateway technology (ThermoFisher, cat #11791043). The CFP-FRB-LAMP1 vector (Willett et al. 2017)was a gift from Rosa Puertollano (NIH). GFP-FRB-EHD1 was kindly provided by Tsukasa Okiyoneda (Kwansei Gakuin University). CFP-FRB-PM (Addgene plasmid #67517) (Varnai et al. 2006), iRFP-FRB-RAB7 (Addgene plasmid #51613), and iRFP-FRB-RAB5 (Addgene plasmid #51612) plasmids (Hammond, Machner, and Balla 2014) were a gift from Tamas Balla (NIH). The mCherry-ARL8B and 2xMyc-SKIP plasmids were a generous gift from Juan S. Bonifacino (Keren-Kaplan and Bonifacino 2021). The pLJC5-LAMP1-RFP-2xFLAG plasmid (Addgene plasmid #102931) (Abu-Remaileh et al. 2017) was a gift from David Sabatini. The EGFP-RILP plasmid (Addgene plasmid #110498) (Mainou and Dermody 2012) was a gift from Terence Dermody.

### Transfection

Transient transfections of HEK293FT cells were performed using Lipofectamine 2000 in Gibco’s Opti-MEM (ThermoFisher, cat #31985088) and incubated for 24 hours prior to fixation or lysis. For siRNA knockdown experiments, siRNA was transfected using Lipofectamine RNAiMAX (ThermoFisher, cat #13778075) and cells were incubated for 48 hours prior to fixation. Transfection of primary astrocytes were performed using Lipofectamine Stem Reagent (ThermoFisher, cat #STEM00015) and incubated for 48 hours before fixation.

### Antibodies

The following primary antibodies were used for immunocytochemistry and Western blot experiments: mouse anti-FLAG M2 (Sigma-Aldrich, cat #F3165, 1:500 for ICC and 1:10,000 for WB), mouse anti-Myc (Roche, cat #C755B26, 1:500 for ICC), mouse anti-LAMP1 (DSHB, cat #H4B3, 1:100 for ICC), rat anti-LAMP2 (DSHB, cat #H4B4, 1:100 for ICC), rat anti-FLAG (Biolegend, cat #637302, 1:200 for ICC), rabbit anti-RAB10 (Abcam, cat #ab237703, 1:2,000 for WB, 1:100 for ICC), rabbit anti-RAB10 (phospho-T73) (Abcam, cat #ab23026, 1:100 for ICC and 1:2,000 for WB), rabbit anti-RAB12 (Proteintech, cat #18843-1-AP, 1:1,000 for WB), rabbit anti-RAB12 (phospho-S106) (Abcam, cat #ab256487, 1:100 for ICC and 1:2,000 for WB), rabbit anti-LRRK2 (Abcam, cat #ab133474, 1:2,000 for WB), rabbit anti-LRRK2 (phospho S1292) (Abcam, cat #ab203181, 1:2,000 for WB), mouse anti-GFP (Roche, cat #11814460001, 1:10,000 for WB), rabbit anti-EEA1 (Cell Signaling Technology, cat #3288, 1:100 for ICC), mouse anti-EEA1 (BD Transduction Laboratories, cat #610457,1:300 for ICC), sheep anti-TGN46 (Biorad, cat #AHP500GT, 1:500 for ICC), rabbit anti-Rab8a (Cell Signaling Technology, cat #6975, 1:500 for ICC), rabbit anti-cyclophilin B (Abcam, cat #ab16045, 1:2,000 for WB), rabbit anti-JIP4 (Cell Signaling Technology, cat#5519, 1:100 for ICC). For staining of the plasma membrane, Phalloidin was used at 1:20 concentration to visualize F-actin (ThermoFisher, cat #A30107).

For ICC, unless otherwise stated, the secondary antibodies were purchased from ThermoFisher. The following secondary antibodies were used: donkey anti-mouse Alexa-Fluor 568 (cat #A10037, 1:500), donkey anti-rabbit Alexa-Fluor 488 (cat #A-21206, 1:500), donkey anti-mouse Alexa-Fluor 488 (cat #A-21202, 1:500), donkey anti-rat Alexa-Fluor 488 (cat #A-21208, 1:500), donkey anti-goat Alexa-Fluor 488 (cat #A-11055, 1:500), donkey anti-rabbit Alexa-Fluor 568 (cat #A10042, 1:500), donkey anti-mouse Alexa-Fluor 647 (cat #A-31571, 1:300), goat anti-rat Alexa-Fluor 647 (cat #A-21247, 1:300).

For WB, all secondary antibodies were used at 1:10,000 dilution: IRDyes 800CW Goat anti-Rabbit IgG (Licor, cat #926–32211) and 680RD Goat anti-Mouse IgG (Licor, cat #926-68070). All blots presented within each figure panel were derived from the same experiment and processed in parallel.

### Immunostaining

HEK293FT cells or mouse primary astrocytes were fixed with 4% PFA for 10 mins, permeabilized with PBS/ 0.1% Triton for 10 minutes and blocked with 5% Donkey Serum (Sigma, cat #D9663) for 1 hour at RT. Primary antibodies were diluted in blocking buffer (PBS supplemented with 1% Donkey Serum) and incubated overnight at 4°C. After three, 5-minute washes with PBS/ 0.1% Triton, secondary fluorescently labeled antibodies were diluted in blocking buffer (PBS supplemented with 1% Donkey Serum) and incubated for 1 hour at RT. Coverslips were washed twice with 1x PBS and an additional 2x with dH_2_O, and mounted with ProLong^®^ Gold antifade reagent (ThermoFisher, cat #P10144).

### SDS PAGE and Western Blotting

Proteins were resolved on 4–20% Criterion TGX precast gels (Biorad, cat #5671095) running at 200V for 40 minutes. Gels were then transferred to nitrocellulose membranes (Biorad, cat #170415) by semi-dry trans-Blot Turbo transfer system (Biorad). The membranes were blocked with Odyssey Blocking Buffer (Licor, cat #927-40000) and then incubated overnight at 4°C with primary antibodies. Afterwards, membranes were washed in TBST (3×5 min) followed by incubation for 1 hour at RT with fluorescently conjugated secondary antibodies as previously stated above. The blots were washed in TBST (3×5 min) and scanned on an ODYSSEY^®^ CLx. Quantitation of bands were performed using Image Studio (Licor).

### Confocal microscopy and analyses

Confocal images were taken using a Zeiss LSM 880 microscope equipped with a 63X 1.4 NA objective. Super-resolution imaging was performed using the Airyscan mode. Raw data were processed using Airycan processing in ‘auto strength’ mode with Zen Black software version 2.3. Only low plasmid expression cells without obvious overexpression artifacts were imaged. For measuring colocalization, Fiji plugin JACoP was used in which Mander’s correlation corrected for threshold was used to quantify LRRK2:pRab colocalization (Bolte and Cordelières 2006). All analyses of colocalization were performed on super-resolution confocal microscopy images. For measuring signal intensity, integrated density of each cell was measured using Fiji with thresholding (ImageJ, NIH). For measuring peripheral vs total lysosomal count, the Spot Counter plugin was used in Fiji. For measuring distance between lysosomes and the nucleus, the surface and spot rendering tools were used on the Imaris software version 9.7.2.

### Statistical analysis

Unpaired Student’s t-tests were used for statistical analyses of experiments with two treatment groups. For more than two groups, we used one-way ANOVA or two-way ANOVA where there were two factors in the model. Tukey’s *post-hoc* test was used to determine statistical significance for individual comparisons in those cases where the underlying ANOVA was statistically significant and where all groups’ means were compared to every other mean. Unless otherwise stated, graphed data are presented as means ± SD. Comparisons were considered statistically significant where *p* < 0.05. *, *p* < 0.05; **, *p* < 0.01; ***, *p* < 0.001; ****, *p* < 0.0001.

## ACKNOWLEDGEMENTS

This research was supported by the Intramural Research Program of the NIH, National Institute on Aging (MRC) and the University of Reading. We thank every member from the Cookson lab for critical feedback. We also thank Rosa Puertollano for kindly providing the CFP-FRB-LAMP1 construct and Juan S. Bonifacino for the gift of the mCherry-ARL8B and 2xMyc-SKIP constructs.

## AUTHOR CONTRIBUTIONS

Conceptualization, L.B-P., M.R.C., J.H.K.; Methodology, L.B-P., J.H.K., A.B.; Formal Analysis, J.H.K., L.B-P.; Investigation, J.H.K., L.B-P.; Writing, J.H.K., L.B-P., M.R.C.; Supervision, L.B-P., M.R.C., P.A.L.; Funding, M.R.C., P.A.L.

## DECLARATION OF INTERESTS

The authors have no competing interests related to this work.

---

**Supplementary Fig 1. Colocalization of endogenous lysosomal and early endosomal markers validate LYSO- and EE-LRRK2 chimeras, respectively.**

To validate the accuracy of both LYSO- and EE-LRRK2 chimeras in translocating LRRK2 to the lysosomal and early endosomal membranes, various resident proteins were stained and colocalization was measured using Mander’s coefficient. For LYSO-LRRK2 (magenta), proteins LAMP2, LAMTOR4, and Cathepsin D (yellow) were measured (A-C). For EE-LRRK2 (magenta), proteins EEA1 and VPS35 (yellow) were measured (D-E). Scale bar= 10μm. (C, E): n=13 cells; SD error bars shown.

**Supplementary Fig 2. The FKBP/FRB complex can be used to trap and activate LRRK2 at various membranes.**

Representative confocal images, Western blots, and the analyses are shown for additional traps using iRFP-FRB-Rab7 (A-E), GFP-FRB-Giantin (F-J), GFP-FRB-EHD1 (K-O), and CFP-FRB-PM (P-T) plasmids, translocating LRRK2 to late endosomes, Golgi, recycling endosomes, and plasma membrane in the presence of rapamycin, respectively. The construct designs for each FRB-trap are illustrated above each representative confocal image (A,F,K,P). All traps are shown in cyan and LRRK2 in magenta. LAMP1, TGN46, Rab8a, and F-actin were chosen as endogenous resident protein markers for each respective membrane (yellow). Western blot analyses of each trap show significant increases in pS1292, pT73 Rab10, and pS106 Rab12 levels when cells are treated with 200 nM rapamycin compared to control cells, and this increase in phosphorylation is abolished after 90 minutes of 1 mM MLi-2 treatment (B-E, G-J, L-O, Q-T). Scale bar= 10 μm. (C-E, H-J, M-O, R-T): one-way ANOVA with Tukey’s post hoc; n=3; SD error bars shown. (C) *F*(2, 6) = 22.40, (D) *F*(2, 6) = 691.5, (E) *F*(2, 6) = 48.89, (H) *F*(2, 6) = 116.4, (I) *F*(2, 6) = 81.12, (J) *F*(2, 6) = 376.3, (M) *F*(2, 6) = 104.6, (N) *F*(2, 6) = 110.5, (O) *F*(2, 6) = 48.55, (R) *F*(2, 6) = 119.8, (S) *F*(2, 6) = 277.7, (T) *F*(2, 6) = 154.4. A schematic illustrating the mechanism in which the LRRK2-FKBP/FRB-membrane trap complex is formed in the presence of rapamycin (U).

**Supplementary Fig 3. Recruitment of Rab10 is limited to a subset of perinuclear, LRRK2-positive lysosomes rather than a limitation on the phosphorylation of Rab10.**

HEK293FT cells expressing the LYSO-LRRK2 chimera (magenta) were stained with total Rab10 (cyan) (A) and cells expressing wildtype LRRK2 are shown as a comparison, with minimal total Rab10 signal (B). Additionally, JIP4 staining (cyan) results in a similar recruitment to lysosomes expressing LYSO-LRRK2 (C) compared to wildtype LRRK2 (D). Overexpression of ARL8B (yellow) and SKIP (not shown) resulted in no total Rab10 recruitment to peripherally situated LRRK2-positive lysosomes (E). Scale bar= 10μm, 15μm.

**Supplementary Fig 4. The centrosome is situated adjacent to pRab10-positive lysosomes.**

Representative confocal images of HEK293FT and U2OS cells stained for LYSO-LRRK2 (magenta), pRab10 (cyan), Pericentrin (green), and RILP (yellow) (A-B). Scale bar= 10μm.

**Supplementary Fig 5. Galectin-3 is observed at peripheral lysosomes after LLOMe treatment in cells expressing ARL8B and SKIP plasmids.**

Representative Airyscan images show GFP-Galectin-3 protein (cyan) is recruited to lysosomes (determined as LAMP2-positive organelles, magenta) in LLOMe-treated cells where ARL8B (yellow) and SKIP (not shown) are expressed. Scale bar= 15 μm, 20 μm.

## Notes

### Competing Interest Statement

The authors have declared no competing interest.

