## Supplementary figures and images for "Lysosomal positioning regulates Rab10 phosphorylation at LRRK2-positive lysosomes"

### Fig S1

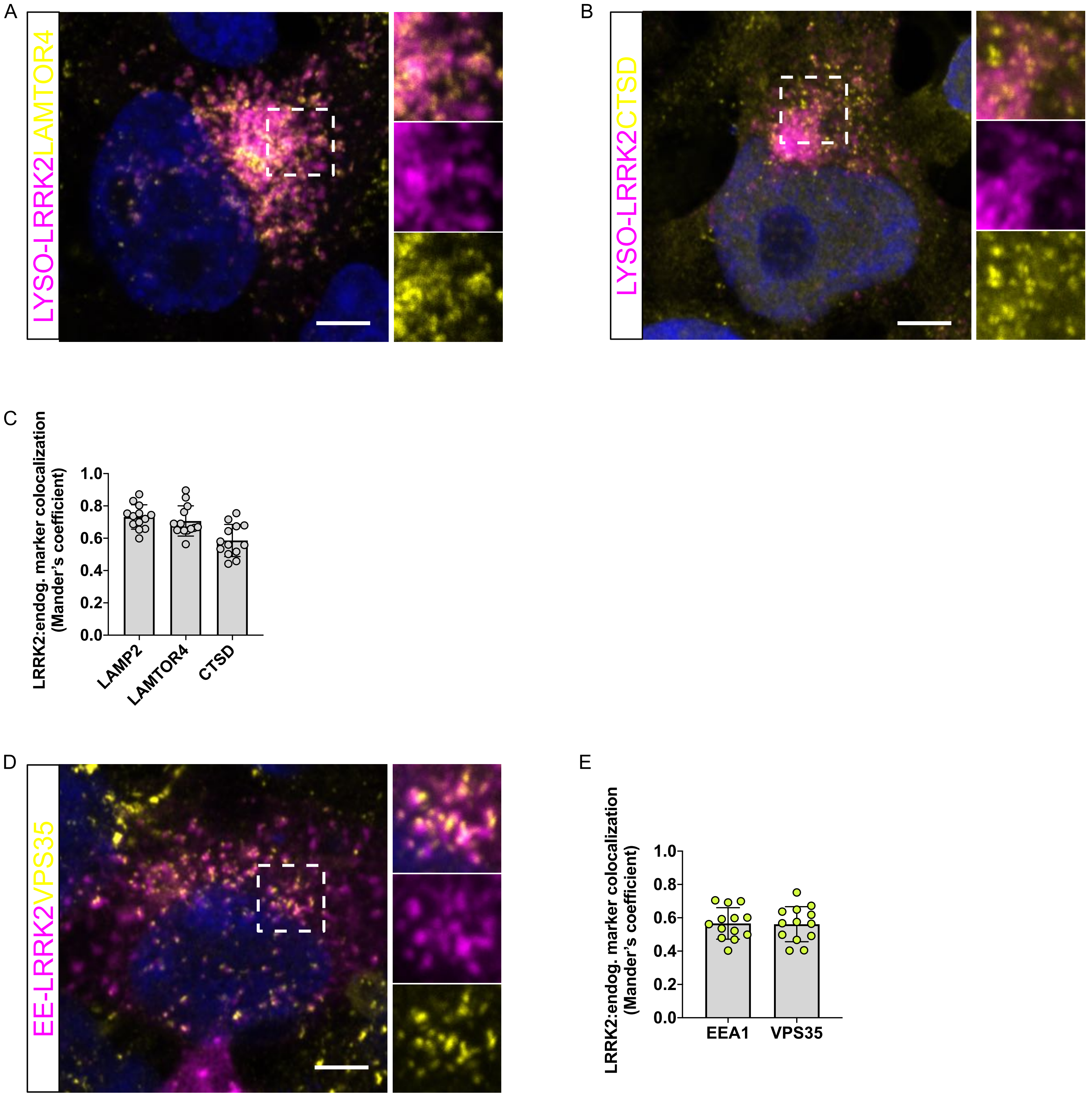


Supplementary Fig 1

### Fig S2

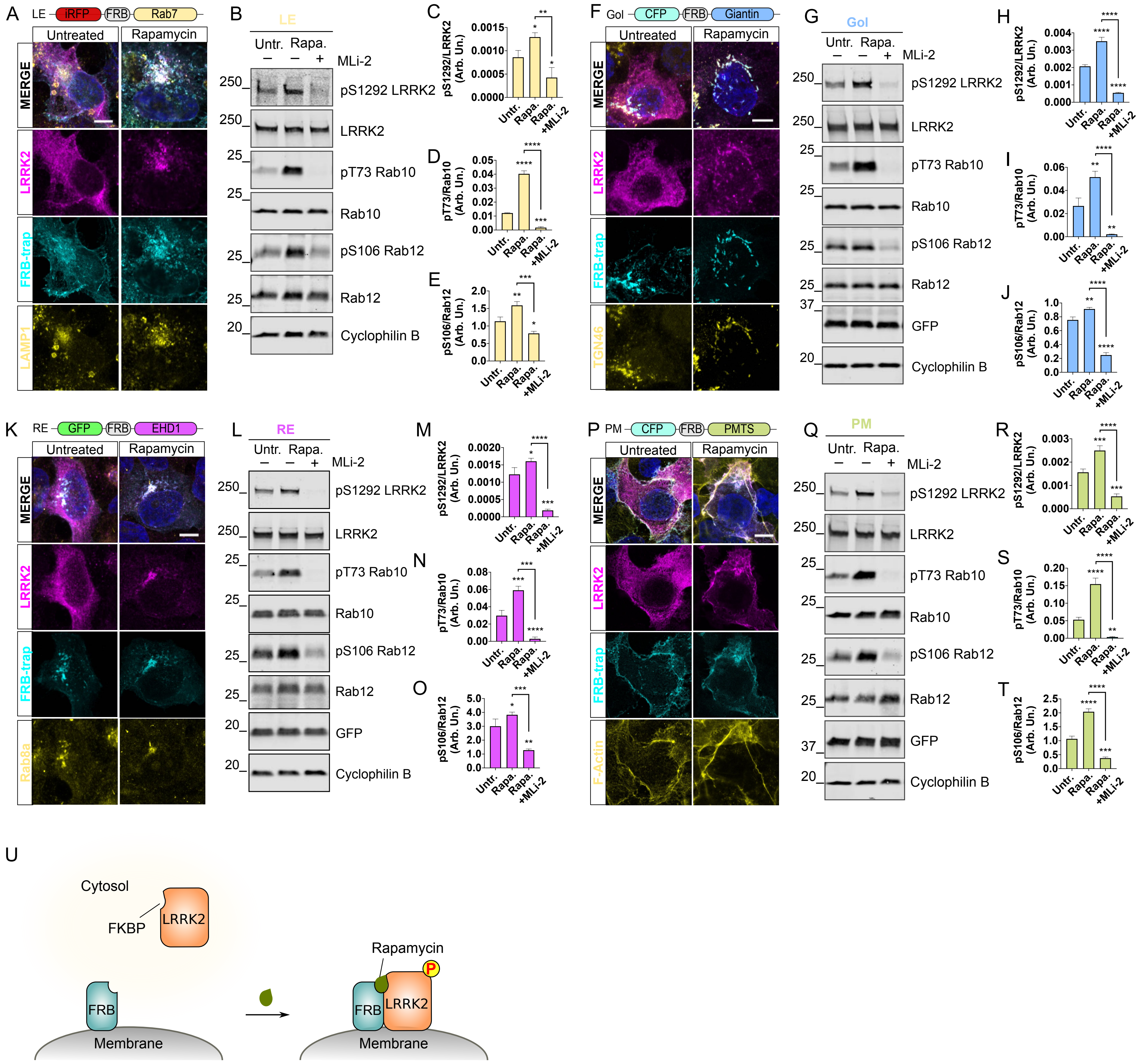


Supplementary Fig 2

### Fig S3

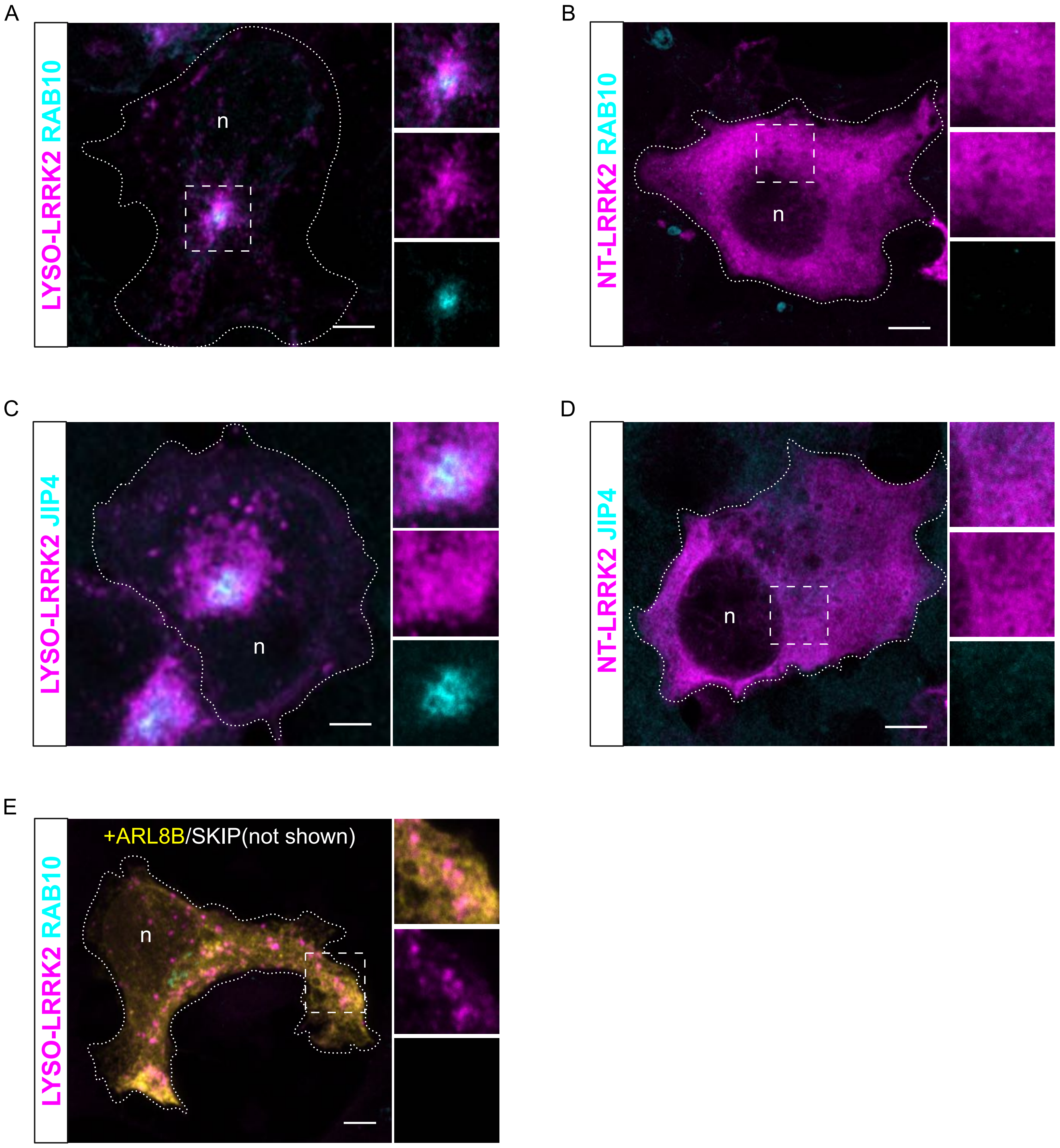


Supplementary Fig 3

### Fig S4

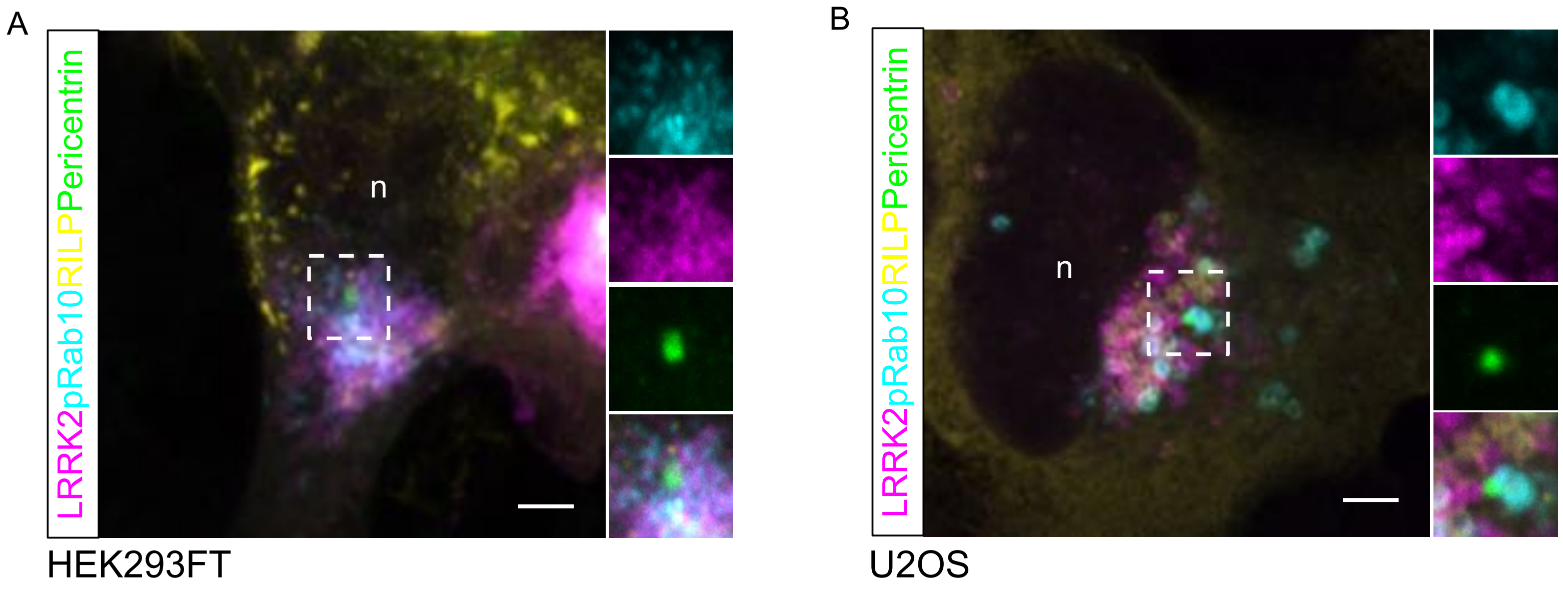


Supplementary Fig 4

### Fig S5

A

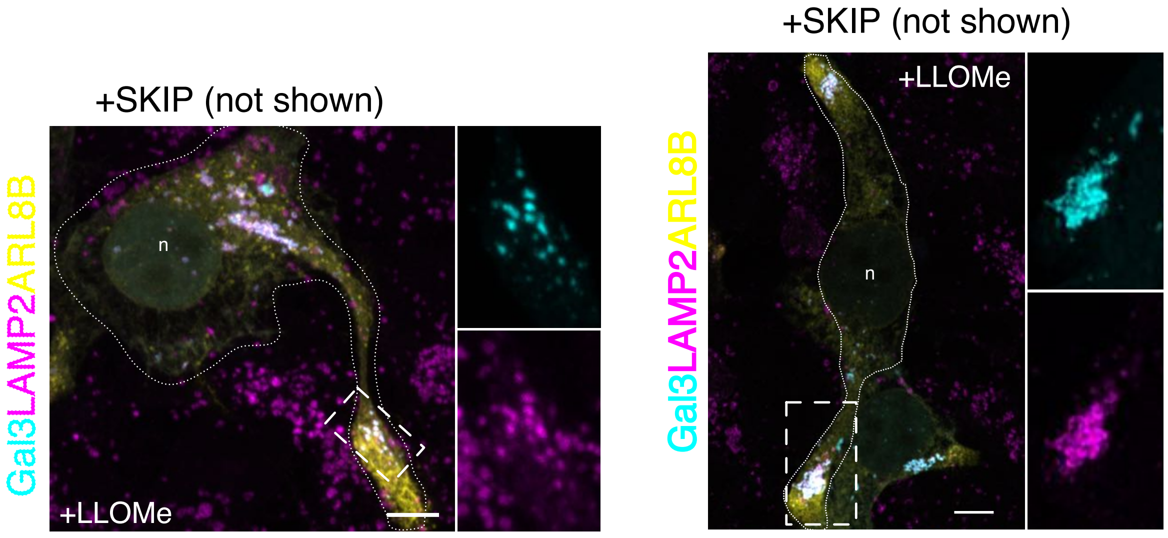


Supplementary Fig 5
